# Cooperating H3N2 influenza virus variants are not detectable in primary clinical samples

**DOI:** 10.1101/223677

**Authors:** Katherine S. Xue, Alexander L. Greninger, Ailyn Pérez-Osorio, Jesse D. Bloom

## Abstract

The high mutation rates of RNA viruses lead to rapid genetic diversification, which can enable cooperative interactions between variants in a viral population. We previously described two distinct variants of H3N2 influenza virus that cooperate in cell culture. These variants differ by a single mutation, D151G, in the neuraminidase protein. The D151G mutation reaches a stable frequency of about 50% when virus is passaged in cell culture. However, it is unclear whether selection for the cooperative benefits of D151G is a cell-culture phenomenon, or whether the mutation is also sometimes present at appreciable frequency in virus populations sampled directly from infected humans. Prior work has not detected D151G in unpassaged clinical samples, but these studies have used methods like Sanger sequencing and pyrosequencing that are relatively insensitive to low-frequency variation. We identified nine samples of human H3N2 influenza collected between 2013 to 2015 in which Sanger sequencing had detected a high frequency of the D151G mutation following one to three passages in cell culture. We deep-sequenced the unpassaged clinical samples to identify low-frequency viral variants. The frequency of D151G did not exceed the frequency of library preparation and sequencing errors in any of the sequenced samples. We conclude that passage in cell culture is primarily responsible for the frequent observations of D151G in recent H3N2 influenza strains.

**IMPORTANCE:** Viruses mutate rapidly, and recent studies of RNA viruses have shown that related viral variants can sometimes cooperate to improve each other’s growth. We previously described two variants of H3N2 influenza virus that cooperate in cell culture. The mutation responsible for cooperation is often observed when human samples of influenza virus are grown in the lab before sequencing, but it is unclear whether the mutation also exists in human infections or is exclusively the result of lab passage. We identified nine human isolates of influenza that had developed the cooperating mutation after being grown in the lab, and performed highly sensitive deep-sequencing of the unpassaged clinical samples to determine whether the mutation existed in the original human infections. We found no evidence of the cooperating mutation in the unpassaged samples, suggesting that the cooperation primarily arises in laboratory conditions.

## INTRODUCTION

RNA viruses like influenza mutate rapidly to form genetically diverse quasispecies. Several recent studies have suggested that interactions between different variants in a quasispecies can promote overall population fitness. In poliovirus, variants generated through spontaneous mutation are important for neurotropism, innate immune suppression, and overall pathogenesis in mouse models (1–3). Other groups have identified cooperative interactions in measles virus (4), West Nile virus (5), hepatitis B virus (6), and Coxsackie virus (7). These cooperative interactions have primarily been observed in cell culture or animal models rather than clinical infections.

We previously described two distinct variants of H3N2 influenza virus that cooperate in cell culture (8). The two variants differ by a single mutation at amino acid 151 of neuraminidase (NA), the protein that releases new virions from host cells. The D151 viral variant, typically encoded as `GAT`, predominates among clinical influenza samples, and it grows robustly in cell culture. The G151 viral variant, typically encoded as `GGT`, binds sialic-acid receptors rather than cleaving them (9, 10) and grows extremely poorly in isolation. However, a mixed population of D151 and G151 viral variants outgrows either single variant in cell culture.

An important question is whether cooperation between these two viral variants is purely a cell-culture phenomenon, or whether the D151 and G151 variants co-exist in natural infections. The D151G mutation is frequently observed when influenza virus is passaged through cell culture (9, 11–16), but it remains unclear whether the G151 variant exists within natural human infections or is primarily a cell-culture artifact. Prior groups that have performed matched clinical sequencing of unpassaged and passaged clinical samples have failed to detect the G151 variant before passaging (13, 15), but these studies have used methods like Sanger sequencing and pyrosequencing that are relatively insensitive to rare variation. More sensitive characterization of clinical samples that give rise to D151G upon lab passage can determine whether this mutation reaches high frequencies in cell culture because it is amplified from low- to modest-frequency standing diversity, or whether it arises spontaneously in the lab.

We sought to determine whether the D151G mutation is present in viral populations isolated from natural human infections. We identified nine clinical samples that, based on prior Sanger sequencing, consisted of a mixture of D151 and G151 viruses after passage in cell culture. We deep-sequenced the original unpassaged nasal swab samples to survey the variation present prior to laboratory growth. The D151G mutation did not exceed the frequency of library preparation and sequencing errors in any of these samples. These results suggest that most variation observed at site 151 results from passage in cell culture rather than standing variation in human infections.

## RESULTS

Most influenza-virus sequences in public databases are determined by Sanger sequencing of clinical isolates that have been passaged one or more times in cell culture (17). A substantial number of recent human H3N2 influenza virus sequences in these databases contain an ambiguous nucleotide at NA site 151 because the lab-passaged samples often converge to a mix of the D151 and G151 variants (8). We sought to compare passaged samples that contained this ambiguous nucleotide at site 151 to unpassaged samples from the same viral infections. We first identified strains from western Washington state in the GISAID EpiFlu database (18) for which Sanger sequencing had reported an ambiguous nucleotide at NA site 151 corresponding to a mix of the D151 and G151 variants (8). Based on the annotations available in the GISAID EpiFlu database, most of these strains had been passaged in cell culture prior to Sanger sequencing.

We obtained original, unpassaged nasal swab samples for the nine strains in Table 1 that contained a mixture of D151 and G151 variants after passage in cell culture. These samples had been collected between 2013 and 2015 and had undergone one to three passages in cell culture prior to sequencing. We performed whole-genome sequencing of the influenza genome from the unpassaged clinical samples using influenza-specific reverse transcription and PCR (19). For each sample, we prepared sequencing libraries in duplicate, beginning from separate reverse-transcription reactions (20). We sequenced each viral sample to an average sequencing depth of 100x-10,000x (Figure 1), allowing us to observe viral variants at frequencies below the limit of detection of Sanger sequencing or pyrosequencing.

**Figure 1:**
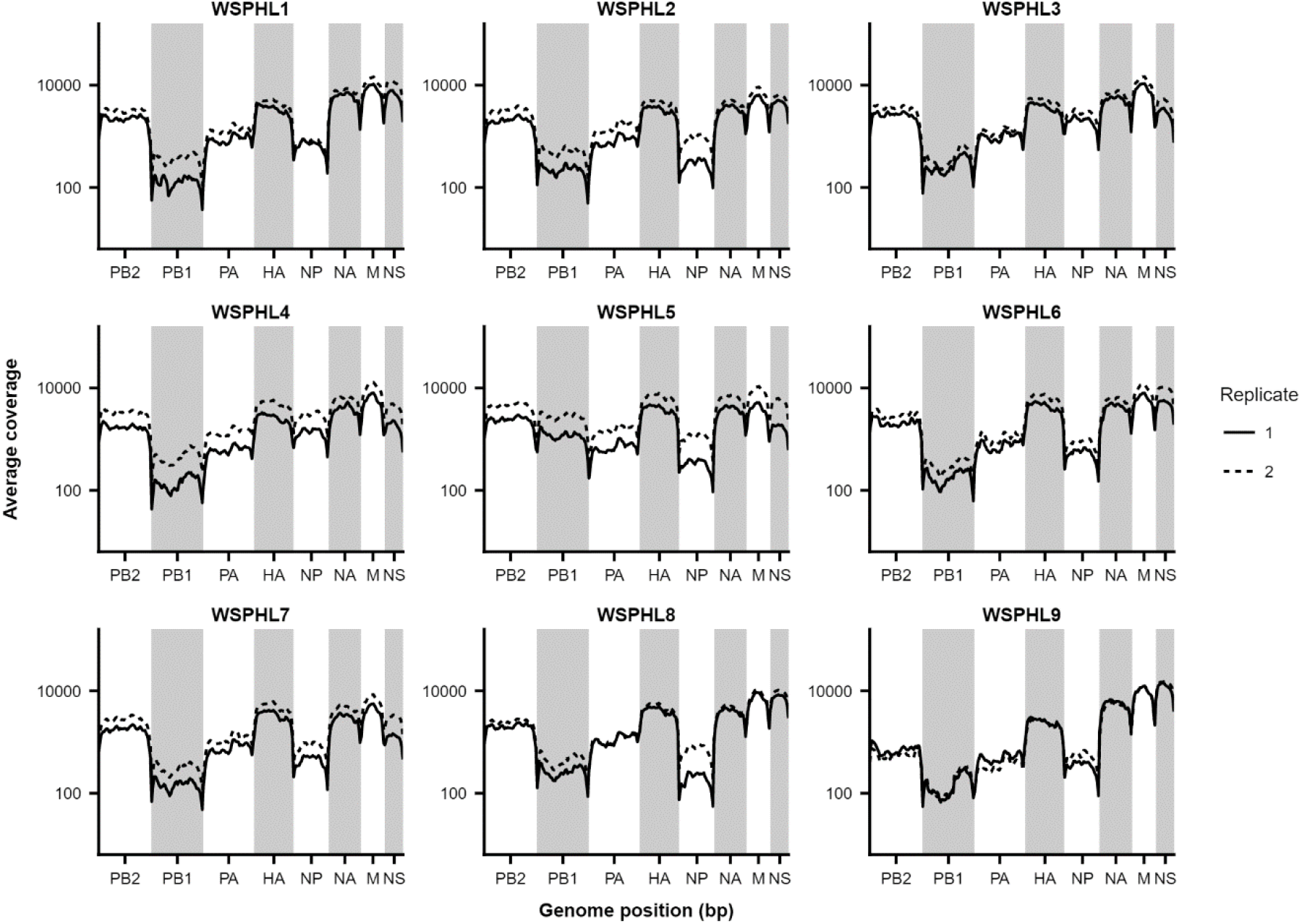
Sequencing coverage along the influenza genome. Average sequencing coverage is plotted for 50 bp bins across the genome, with library replicates shown in solid and dashed lines.

**Table 1.** Strains deep-sequenced in this study. Genotypes were determined through Sanger sequencing of passaged isolates, and are taken from those reported in the GISAID EpiFlu database. Annotations of passage history are not standardized, but C[N] generally refers to N passages of the virus in cell culture prior to sequencing, and S[N] generally refers to N passages of the virus in MDCK-SIAT1 cells (17). For the genotype at site 151, an annotation of X indicates a mix of D151 and G151 in the original Sanger sequencing. The Ct value is the amount of viral material in the original clinical sample as determined by qPCR.

| Sample | Strain | Passage history | Genotype, site 151 | Ct Value |
| --- | --- | --- | --- | --- |
| WSPHL1 | A/Washington/10/2013 | C1/C1 | X | 23.19 |
| WSPHL2 | A/Washington/13/2013 | C1 | X | 17 |
| WSPHL3 | A/Washington/17/2013 | C2 | X | 24.57 |
| WSPHL4 | A/Washington/18/2013 | C3 | X | 21.52 |
| WSPHL5 | A/Washington/08/2014 | C1 | X | 22.8 |
| WSPHL6 | A/Washington/07/2015 | S3 | X | 23.78 |
| WSPHL7 | A/Washington/24/2015 | S3 | X | 17.4 |
| WSPHL8 | A/Washington/32/2015 | S3 | X | 18.69 |
| WSPHL9 | A/Washington/36/2015 | S3 | X | 25.03 |

We identified all minor viral variants present at a frequency of at least 3% in the viral genome in both library replicates (Table 2). We did not observe the D151G variant in any of the nine clinical samples under these variant-calling criteria. To ensure that we were not missing extremely low-frequency variation, we calculated the frequency of D151G in each clinical sample based on the frequency of G-to-A mutations at the second nucleotide position of NA site 151. We compared this frequency to the frequency of G-to-A mutations at other sites across the genome (Figure 2). Minor-variant frequencies at NA site 151 fell well within the range of error expected through library preparation and sequencing errors. Therefore, we conclude that the D151G variant was not present at appreciable frequencies in the original clinical infections. Instead, the mutation must have arisen *de novo* or been enriched from an extremely low frequency during passage in cell culture.

**Table 2.**
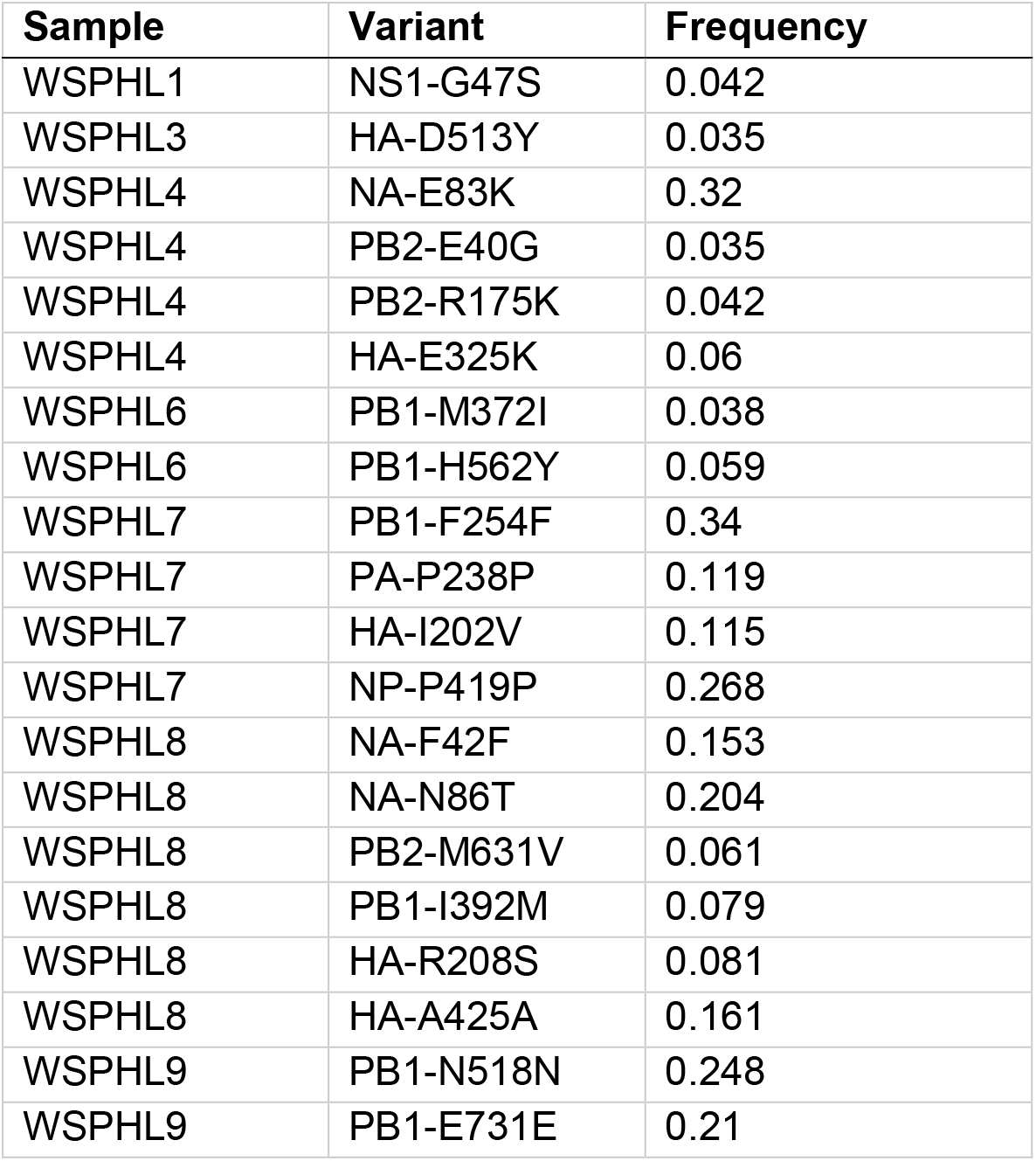
Within-host variants identified through deep sequencing. Sites were called as variable if a non-consensus base exceeded a frequency of 0.03, given a sequencing coverage of at least 100x, in both sequencing replicates.

**Figure 2:**
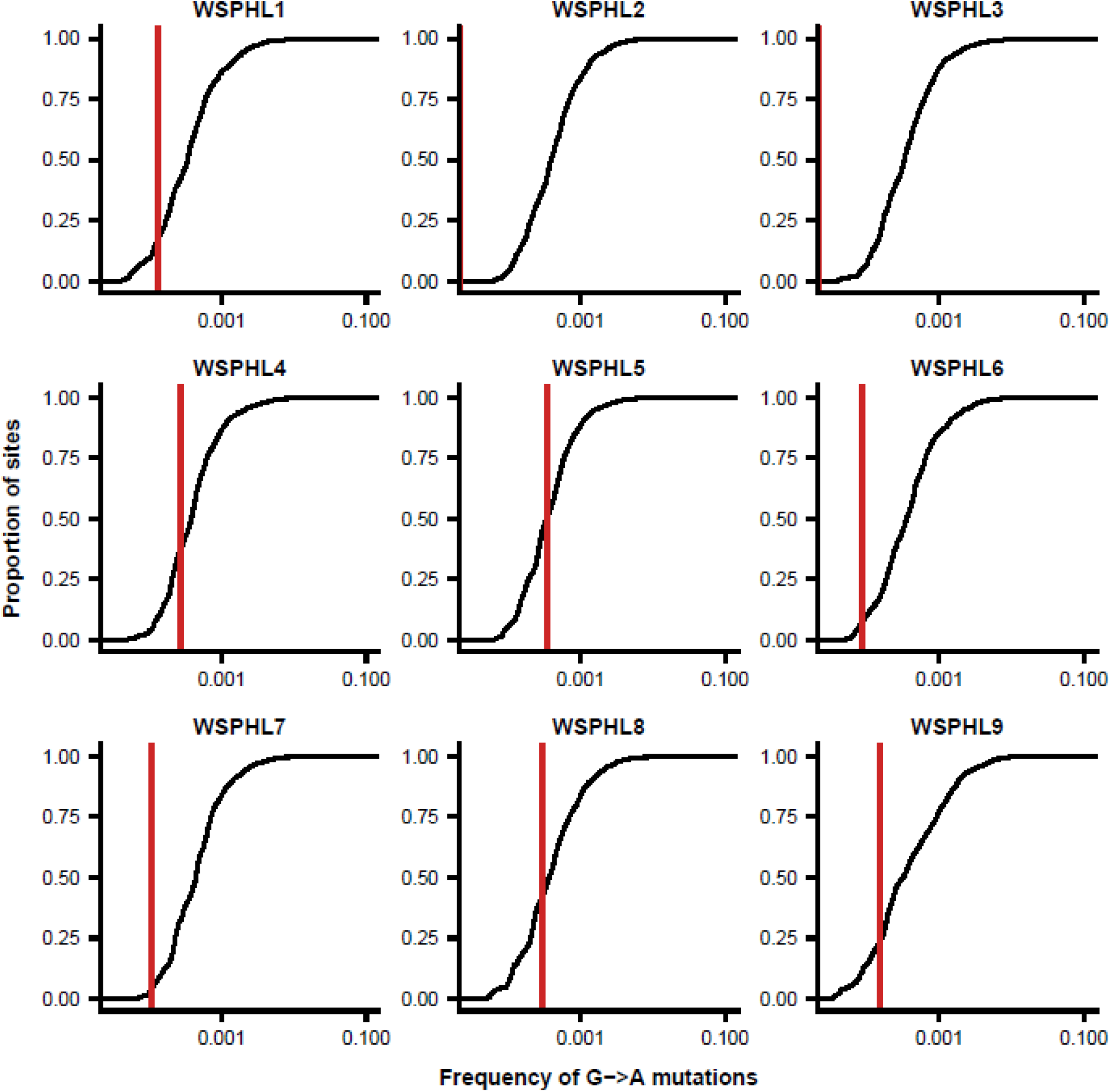
D151G does not exceed the frequency of library preparation and sequencing errors in unpassaged clinical samples. Shown is the distribution of frequencies of G-to-A mutations across the genome for each clinical sample. Typically, the D151 viral variant is encoded by the nucleotides `GAT`, and the G151 variant is encoded as `GGT`, meaning that D151G arises as the result of a G-to-A mutation. The red vertical line shows the proportion of G-to-A mutations at codon position 2 of amino-acid site 151 of NA, which corresponds to the frequency of D151G. In cases where no G-to-A mutations were identified at this site, this red line is not shown. At each nucleotide site in the genome with consensus identity G, we calculated the total proportion of reads reporting an identity of A at that site and averaged this proportion between both replicate libraries. As expected, G-to-A mutations make up less than 0.1% of total sequencing reads at most sites in the genome and are probably errors introduced through library preparation and sequencing.

## DISCUSSION

The results of our deep-sequencing study support prior studies that failed to detect the D151G mutation in unpassaged clinical samples using Sanger sequencing or pyrosequencing methods (13, 15). In the GISAID EpiFlu database, mixed populations of D151 and G151 viral variants are common in clinical samples that have been passaged in cell culture, but these mixed populations are rare among unpassaged and egg-passaged populations (8). It is impossible to rule out the possibility that the D151G mutation reaches appreciable frequencies in some natural human infections, but strong and repeated selection for cooperation in cell culture seems to account for its prevalence among sequences in public databases.

It is interesting to speculate about what biological factors might cause a variant that is rare in natural human infections to be strongly selected in cell culture. Influenza strains often acquire stereotypical mutations when they are grown in eggs (21, 22), but these passage adaptations appear to be less common in cell culture, particularly for `MdCk-SIAT`1 cells (17, 23). Nevertheless, differences in the types and distributions of cell-surface receptors between MDCK-SIAT1 cells and human airways could account for some of the differences in genotypes we observe at NA site 151.

We also previously observed that cooperation is stronger at high multiplicities of infection (MOI) (8). Viral load can be high during natural infections (Table 1), but recent studies of natural human infections have found that the effective reassortment rate is limited, suggesting that spatial heterogeneity within the host may limit viral circulation and co-infection (24). Moreover, human influenza infections, as well as those in animal models (25), experience a severe transmission bottleneck that greatly limits the genetic diversity initially present in an infection (26–28). In contrast, viral populations can rapidly reach high MOIs in cell culture (29). These different growth conditions may also promote the emergence of D151G within cell culture, but not natural infections.

Our study also underscores the importance of sequencing directly from unpassaged clinical samples. Mutations like D151G accumulate in cell culture within just a few passages and affect downstream analyses like inferences of positive selection (17). Careful records of passage histories combined with deep-sequencing of unpassaged clinical samples can help distinguish natural variation from that generated in the lab.

## MATERIALS AND METHODS

### Viral samples

We downloaded the set of 66 sequences in the Global Initiative on Sharing All Influenza Data (GISAID) EpiFlu database (18) corresponding to all full-length NA coding regions from human H3N2 influenza A isolates collected from January 1, 2000 to August 26, 2015 and submitted from Seattle, Washington, or Shoreline, Washington (**Table S1**). We pairwise aligned each sequence to the A/Hanoi/Q118/2007 (H3N2) coding sequence (Genbank accession CY104446) using the program needle from EMBOSS version 6.6.0 (30). For each sequence, we determined the genotype at site 151 and assigned the genotype X if there was an ambiguous nucleotide at that site. We identified sequences with ambiguous identities at site 151, suggesting the presence of mixed viral populations, and we extracted passage histories based on the metadata available in the GISAID Epiflu database. For the nine strains described in Table 1, we were able to obtain aliquots of the original, unpassaged nasal swab samples in viral transport media.

### Viral deep sequencing

We performed viral deep sequencing as previously described (19). In brief, we extracted viral RNA from unpassaged clinical samples using the QIAamp Viral RNA Mini Kit (Qiagen) according to manufacturer’s instructions. We reverse-transcribed the viral RNA using the Superscript III First-Strand Reaction Mix (Thermo Fisher) and an equimolar mix of the influenza-specific primers 5′-TATTGGTCTCAGGGAGCAAAAGCAGG-3′ and 5′-TATTGGTCTCAGGGAGCGAAAGCAGG-3′, which both bind to the conserved U12 region at one end of each influenza gene. The two primers differ by a single nucleotide to account for a known polymorphism in the region. We incubated the reverse-transcription reactions at 25 degrees C for 10 minutes (to help the short primer anneal), 50 degrees C for 50 minutes, and 85 degrees C for 5 minutes. We amplified the influenza genome using a mixture of 24 primers that bind to the ends of each influenza gene (31). For each gene, one primer binds to the conserved U13 region at one end of the gene, and two primers bind to the conserved U12 region at the other end of the gene, allowing for the known polymorphism in the U12 region. We performed 35 cycles of PCR using an annealing temperature of 55 degrees C and an extension time of 3 minutes. We purified the PCR product using 1X AMPure beads (Beckman Coulter) and prepared libraries for Illumina sequencing using Nextera XT (Illumina) tagmentation. We sequenced the libraries on a NextSeq 500 platform (Illumina) with 150 bp paired-end reads. We performed all library preparation and sequencing in duplicate, starting from independent reverse-transcription reactions (20).

### Analysis of deep-sequencing data

We first used bowtie2 (32) to filter out reads that mapped to the human genome. Remaining reads are available in the SRA as BioProject PRJNA412675. We trimmed adapters from the raw reads using cutadapt version 1.8.3 (33). We first aligned the reads to the A/Victoria/361/2011 genome using bowtie2 and the – `very-sensitive` setting, then we used custom scripts to generate a new consensus genome sequence for each viral sample. We then re-aligned the reads to the corresponding consensus sequence and removed PCR duplicates using picard version 1.43. We used custom scripts to filter out base calls with a quality score below 20, tally the total number of high-quality bases at each genome position, and annotate each variant’s codon position. We performed these initial analyses separately for each replicate library. We reported only variants that were located in protein-coding sequence.

### A note on codon numbering and gene annotation

We numbered HA codons according to the H3 numbering system. This HA numbering scheme assigns 1 to codon 17 of the full HA gene, which is the beginning of the mature HA protein. The codons for all other genes are numbered sequentially beginning with one at the N-terminal methionine. The M1 and M2 genes have 27 bp of in-frame and 44 bp of out-of-frame overlap, and the NS1 and NEP genes have 30 bp of in-frame and 251 bp of out-of-frame overlap. We annotated variants separately for each gene if they occurred in these regions of overlap.

### Data and code availability

Sequencing reads are available on the SRA as BioProject PRJNA412675. The computer code that performs the analyses is available on Github at https://github.com/ksxue/D151G-clinical-public.

## ACKNOWLEDGEMENTS

This study was funded by R01GM102198 from the NIGMS, R01AI127893 from the NIAID, a Howard Hughes Medical Institute Faculty Scholar Award, and a Simons Foundation Faculty Scholar Award to JDB. KSX was funded by an NSF Graduate Research Fellowship under grant number DGE-1256082 and a graduate fellowship from the Fannie and John Hertz Foundation. The funders had no role in study design, data collection and interpretation, or the decision to submit the work for publication.

